# Towards a standard benchmark for phenotype-driven variant and gene prioritisation algorithms: PhEval - Phenotypic inference Evaluation framework

**DOI:** 10.1101/2024.06.13.598672

**Authors:** Yasemin Bridges, Vinicius de Souza, Katherina G Cortes, Melissa Haendel, Nomi L Harris, Daniel R Korn, Nikolaos M Marinakis, Nicolas Matentzoglu, James A McLaughlin, Christopher J Mungall, Aaron Odell, David Osumi-Sutherland, Peter N Robinson, Damian Smedley, Julius OB Jacobsen

## Abstract

**Background:** Computational approaches to support rare disease diagnosis are challenging to build, requiring the integration of complex data types such as ontologies, gene-to-phenotype associations, and cross-species data into variant and gene prioritisation algorithms (VGPAs). However, the performance of VGPAs has been difficult to measure and is impacted by many factors, for example, ontology structure, annotation completeness or changes to the underlying algorithm. Assertions of the capabilities of VGPAs are often not reproducible, in part because there is no standardised, empirical framework and openly available patient data to assess the efficacy of VGPAs - ultimately hindering the development of effective prioritisation tools.

**Results:** In this paper, we present our benchmarking tool, PhEval, which aims to provide a standardised and empirical framework to evaluate phenotype-driven VGPAs. The inclusion of standardised test corpora and test corpus generation tools in the PhEval suite of tools allows open benchmarking and comparison of methods on standardised data sets.

**Conclusions:** PhEval and the standardised test corpora solve the issues of patient data availability and experimental tooling configuration when benchmarking and comparing rare disease VGPAs. By providing standardised data on patient cohorts from real-world case-reports and controlling the configuration of evaluated VGPAs, PhEval enables transparent, portable, comparable and reproducible benchmarking of VGPAs. As these tools are often a key component of many rare disease diagnostic pipelines, a thorough and standardised method of assessment is essential for improving patient diagnosis and care.

## Background

Rare diseases are defined as diseases affecting fewer than 200,000 individuals in the United States or fewer than 1 in 2,000 individuals in the EU [1]. It is estimated that over 400 million people worldwide are affected by some rare disease [2]. Oftentimes these diseases are so rare and complex that a patient may take years or even decades to receive an accurate diagnosis. Many rare diseases can be caused by extremely small errors in a patient’s genetic code. Accurately identifying genetic variants in patient data can greatly aid in the understanding of their conditions [3]. Since the average human has around 4-5 million variations in their genetic profile, most of which are not relevant to their disease(s), the process of utilising genomic information in diagnosis is twofold: identifying a specific patient’s variants by sequencing and then prioritising those variants to only those with a high likelihood of being relevant to the patient’s phenotypes of interest [4, 5].

Next-generation sequencing (NGS) techniques, such as Whole Exome Sequencing (WES) and Whole Genome Sequencing (WGS), provide fast and cost-effective ways to quickly profile patient genetic information content. The extensive amount of genetic profiles generated by NGS techniques is then processed by variant and gene prioritisation algorithms (VGPAs). These algorithms implement a variety of methodologies, including leveraging phenotype ontologies, integrating cross-species data, analysing gene expression patterns, among others, with specific approaches varying across tools. Over the years, VGPAs have demonstrated utility in clinical diagnostics and research, enabling clinicians and researchers to pinpoint potentially pathogenic variants responsible for rare diseases.

Phenotype data, which describes an individual’s observable traits, clinical features and medical history, plays a significant role in understanding the potential impact of genetic variants. The Human Phenotype Ontology (HPO) is a specially designed vocabulary and organisational structure which enumerates and categorises all human phenotypes [6]. HPO is widely used in diagnostic pipelines in many clinical and research settings.

In rare disease diagnosis, the integration of phenotype data into VGPAs has proven crucial for enhancing the accuracy and clinical relevance of genomic variant interpretation [7]. A key component of the integration of phenotype data and VGPAs is the utilisation of HPO terms, which serves as a crucial link between genomic information and clinical medicine, with the goal of covering phenotypic abnormalities in monogenic diseases [6]. More than 20 VGPA tools leverage HPO to incorporate phenotypic data in their algorithms [8], including well-known examples such as Exomiser [9], LIRICAL [10], and Phen2Gene [11]. Work by Robinson et al., Jacobsen et al., and Thompson et al. has provided insights into improving diagnostic yield, defined as the proportion of cases where a causative entity is correctly identified, achieved by integrating phenotype data within variant prioritisation algorithms [8, 12, 13].

Our group has highlighted the importance of combining variant and phenotype scores into a unified score for the effective prioritisation of genetic variants when utilising Exomiser [9], a VGPA developed by the Monarch Initiative [14]. We demonstrated a significant advancement in the accuracy of Exomiser for predicting relevant disease variants when utilising a combination of genomic and phenotypic information. On a dataset of 4877 patients with a confirmed diagnosis and both genomic and phenotypic information, Exomiser correctly identified the diagnosis as the top-ranking candidate in 82% of cases, compared to only 33% and 55% when solely considering the variant or phenotype scores respectively [8]. The incorporation of phenotype data from both mammalian and non-mammalian organisms into variant prioritisation algorithms has proven to increase accuracy even further. In work by Bone et al. the integration of diverse organism data, encompassing human, mouse, and zebrafish phenotypes, in conjunction with the phenotypes associated with mutations of interacting proteins, led to a substantial improvement of 30% in performance, with 97% of known disease-associated variants from the Human Gene Mutation Database (HGMD) detected as the top-ranked candidate in exomes from the 1000 Genomes project that were spiked with the causal variant [15].

Given variant or gene prioritisation algorithms (VGPAs) are critical diagnostic tools, it is critical that they should be benchmarked before they are utilised in healthcare. Numerous benchmarking studies have been conducted by researchers [15–17], primarily with the objectives of evaluating the performance of new prioritisation algorithms, performing comparative analyses with existing algorithms, and executing comprehensive reviews of algorithms already in use [10, 11, 18]. However, many of these benchmarks face challenges with reproducibility due to insufficient documentation of benchmarking methodologies and closed data sets. Academic software providers and commercial vendors alike often make effectiveness claims, but these are difficult for third parties to independently reproduce and validate. A key step towards enhancing benchmark rigour involves providing transparent and detailed documentation, including data sources and versions, algorithm versions, data pre- processing steps, parameter settings, and provided uniform output from each tool. An anecdote that illustrates the importance of clearly documenting benchmarking methodologies and providing the experimental pipeline in a reproducible manner is demonstrated by a comparative analysis of ten gene-prioritisation algorithms performed by Yuan et al. The study revealed a significant variance in Exomiser’s performance compared to previous benchmarking outcomes [19]. In response, Jacobsen et al. clarified this discrepancy by identifying crucial differences in the parameter settings employed by Yuan et al., which were the likely explanation for the observed differences in the performance of Exomiser [20]. Fortunately, Yuan et al. included the data necessary for reproducing their study, which played a pivotal role in facilitating the correction of discrepancies.

Beyond concerns related to documentation and reproducibility, another significant issue in present benchmarks is the lack of standardisation, particularly concerning the types of test data and performance evaluation metrics used. Consequently, some test datasets may exhibit better performance than others, and the choice of which metrics to use may favour specific algorithms, introducing an element of subjectivity. Such variability raises questions about the consistency and comparability of benchmark results across studies. Additionally, while synthetic datasets are valuable tools for controlled evaluations, they come with inherent limitations in their ability to represent real-world scenarios. Simulated datasets are typically designed based on simplified models, and as such, may not encompass the full spectrum of genetic variants and phenotypic complexities encountered in clinical settings. The use of simulated data may inadvertently favour algorithms that perform well under the specific conditions and assumptions embedded in these datasets, potentially leading to results that diverge significantly from those observed in real-world applications. These challenges collectively create conditions that can make achieving rigour in benchmarking studies more difficult.

There is a notable scarcity of benchmarks specifically designed for evaluating VGPAs. This hinders the objective assessment and comparison of different VGPAs. Without established benchmarks, researchers and clinicians encounter difficulties in gauging the accuracy, efficiency, and clinical relevance of different algorithmic approaches. The absence of these benchmarks also limits opportunities for identifying areas of improvement and innovation within the field of VGPA, potentially slowing down progress in rare disease diagnostics and genomic medicine as a whole.

Benchmarking VGPAs that rely on phenotype data presents a multifaceted challenge due to the extra algorithmic complexity required to monitor performance over time, as well as the additional requirements for test data, pre-processing and analysis. One key hurdle lies in the necessity to preprocess and transform the test data effectively. Patient phenotypic profiles are typically represented as a collection of HPO IDs (an alphanumeric code assigned to each specific phenotype term), and the descriptive label or name associated with each phenotype. However, the divergence in data formats expected by different VGPAs - from simple flat lists to highly structured - complicates the benchmarking process. Here, the GA4GH Phenopacket-schema aims to provide a solution, serving as a standardised and extensible format for representing an individual’s disease and phenotype information, facilitating the consistent exchange of phenotypic data and playing a crucial role in genomics research by aiding in the understanding between genetic variations and observable traits [21].

The complexity of benchmarking these algorithms increases due to the variety of different interfaces that are needed to support these methods. Each algorithm must be individually invoked and executed, often demanding a significant amount of computational resources. Additionally, the logistical aspects of coordinating diverse software components with a variety of implementation details such as programming language and dependencies adds to the complexity. Configuring each VGPA correctly can often involve managing complex configuration files, where performance depends on understanding and fine-tuning multiple parameters. Ensuring all data dependencies, such as paths to data references and resources, are properly set up is also a key part of this challenge. Even more complexity is created by the need to ensure the desired versions of each tool and input data version are used and to execute them in a systematic and reproducible manner.

Beyond the individual execution of algorithms, the benchmarking process also has to harmonise the diverse output formats generated by these tools. To enable meaningful comparisons and evaluations, the outputs must be transformed into a uniform format. This standardisation allows for consistent and structured analysis across different algorithms, ensuring that the results are interpretable and facilitating fair assessments of their performance.

To tackle the absence of standardised benchmarks and data standardisation for VGPAs, we developed PhEval, a novel framework that streamlines the evaluation of VGPAs that incorporate phenotypic data. PhEval is built on the Phenopacket-schema, a GA4GH and ISO standard for sharing detailed phenotypic descriptions with disease, patient, and genetic information, enabling clinicians and other researchers to build and share more complete models of disease in a standardised format. PhEval offers the following value propositions:

- Automated processes: PhEval automates various evaluation tasks, enhancing efficiency and reducing manual effort.
- Standardisation: The framework ensures consistency and comparability in evaluation methodologies, promoting reliable assessments.
- Reproducibility: PhEval facilitates reproducibility in research by providing a standardised platform for evaluation, allowing for consistent validation of algorithms.
- Comprehensive benchmarking: PhEval enables thorough benchmarking of algorithms, allowing for well-founded comparisons and insights into performance.

### Implementation

PhEval is a framework designed to evaluate variant and gene prioritisation algorithms that incorporate phenotypic data to assist in the identification of possibly disease-causing variants. PhEval is specifically designed for evaluating monogenic diseases, where a single causative gene, variant, or disease is expected per case. The framework includes (1) a modular library available on the Python Package Index (PyPI) repository of software that provides a command-line interface (CLI) to handle benchmarks efficiently, (2) an interface for implementing custom VGPA runners as plugins to PhEval, (3) a workflow system for orchestrating experiments and experimental analysis and (4) a set of test corpora.

### PhEval CLI and VGPA runners

The PhEval CLI comprises core and utility commands. The main command is “pheval run”, which executes the variant/gene prioritisation runners. Additionally, there are a collection of utility methods which facilitate some procedures such as generating “noisy” phenopackets to assess the robustness of VGPAs when less relevant or unreliable phenotype data is introduced (see Section on Test Corpora). The “generate-benchmark-stats” command allows users to evaluate and compare the performance of VGPAs algorithms by plotting a graph.

Another core component of PhEval is an extensible system to help enable the execution of a wide variety of VGPAs. To ensure that PhEval can execute and assess VGPA tools in a standardised manner, developers must implement three abstract methods that ensure: (1) the data supplied by the PhEval framework is transformed into whatever input format the VGPA requires, (2) the VGPA tool can be executed using a standardised “run” method (described above, the main entry point for each benchmarking execution) and (3) the data produced by the VGPA tool is converted into the standardised representation required by PhEval to provide a comparison of performance of tools.

To minimise development effort, developers do not need to rewrite or modify their existing tools. Integration should be straightforward for developers familiar with the tool being benchmarked, as they only need to define how it interfaces with PhEval through a tool-specific runner. Developers can use a cookiecutter template to generate a pre-configured project structure and refer to detailed guides on implementing runners and methods.

All processes are described as part of the general PhEval documentation (https://monarch-initiative.github.io/pheval/developing_a_pheval_plugin/), and a concrete reference implementation is available at: (https://github.com/monarch-initiative/pheval.exomiser)

### PhEval experimental pipeline

The PhEval benchmarking process can be broadly divided into three distinct phases: the data preparation phase, the runner phase, and the analysis phase.

*The data preparation phase*, as well as automatically checking the completeness of the disease, gene and variant input data and optionally preparing simulated VCF files if required, gives the user the ability to randomise phenotypic profiles using the PhEval corpus “scramble-phenopackets” command utility, allowing for the assessment of how well VGPAs handle noise and less specific phenotypic profiles when making predictions.

*The runner phase* is structured into three stages: *prepare*, *run*, and *post-process*. While most prioritisation tools are capable of handling some common inputs, such as phenopackets and VCF files, the *prepare* step plays a crucial role in adapting the input data to meet the specific requirements of the tool. For instance, one of the VGPAs we tested, Phen2Gene, which is a phenotype-driven gene prioritisation tool, lacks the ability to process phenopackets during its execution. To address this limitation, the *prepare* step serves as a bridge, facilitating necessary data preprocessing and formatting. In the case of Phen2Gene, potential solutions during the *prepare* step may involve parsing the phenopackets to extract the HPO terms associated with each sample, subsequently providing them to the Phen2Gene client in the run step. Alternatively, it may entail the creation of input text files containing HPO terms that can be processed by Phen2Gene.

In the *run* step, the VGPA is executed, applying the selected algorithm to the prepared data and generating the tool-specific outputs. Within the *run* stage, an essential task is the generation of input command files for the algorithm. These files serve as collections of individual commands, each tailored to run the targeted VGPA on specific samples. These commands are configured with the appropriate inputs, outputs and specific configuration settings, allowing for the automated and efficient processing of large corpora.

Finally, the *post-processing* step takes care of harmonising the tool-specific outputs into standardised PhEval TSV format, ensuring uniformity and ease of analysis of results from all VGPAs. In this context, the tool-specific output is condensed to provide only two essential elements, the entity of interest, which can either be a variant, gene, or disease, and its corresponding score. For example, Exomiser’s JSON output which is rich in content, is parsed to extract only the gene symbols, identifiers and the corresponding score required for the standardised output and subsequent benchmarking. Developers can implement custom parsers in their runners to extract only the relevant data fields required. PhEval then assumes the responsibility of subsequent standardisation processes. This involves the reranking of the results in a uniform manner, based on the original scores provided by the tool, ensuring that fair and comprehensive comparisons can be made between tools. The reranking ensures that all tools are assessed using the same ranking scale, enabling unbiased comparisons regardless of differences in their native ranking mechanisms. PhEval offers an array of utility methods, readily available for integration into plugins by developers, facilitating the seamless generation of standardised PhEval outputs. These methods not only include the generation of the standardised output from extracted essential elements, but also methods for calculating the end position of a variant, converting gene names to specific gene identifiers and vice versa, contributing to an efficient post-processing workflow and ensuring the consistent and standardised representation of the results.

*In the analysis phase*, PhEval generates comprehensive statistical reports based on standardised outputs from the runner phase. This process enables rigorous assessment by comparing these results of VGPAs with the known causative variants (causal variants are provided in the set of phenopackets as an evaluation suite). These reports include both ranked metrics and binary classification. In our evaluation, PhEval uses a rank-based evaluation system, where true positives are defined as the known causative entity ranked at position 1 in the results, false positives are any other entity ranked at position 1, true negatives are any entity ranked at a position other than 1, and false negatives are the known causative entity ranked at a position other than 1. The stringent cutoff of considering only the top ranked entity as a true positive reflects the clinical context of monogenic disease diagnosis, where a single causative entity is expected, and prioritising rank 1 is key for streamlining diagnostic workflows and minimising time-intensive reviews. The ranked metrics we currently support calculating are: the count of known entities found in the top-n ranks, mean reciprocal rank (MRR), precision@k, mean average precision@k, f-beta score @k, and normalised discounted cumulative gain (NDCG) @k. The following binary classification metrics we support are: sensitivity, specificity, precision, negative predictive value, false positive rate, false discovery rate, false negative rate, accuracy, f1-score, and Matthews correlation coefficient. The framework currently offers robust support for analysing and benchmarking prioritisation outcomes related to variants, genes, and diseases, ensuring a thorough evaluation of its performance.

PhEval employs a Makefile (GNU-make) strategy to organise all necessary steps. The GNU-make framework enables easy orchestration of the process of building data corpora, obtaining and installing tools, running VGPAs, and creating benchmarking reports. This ensures that each phase - data preparation, runner, and analysis - is carried out in a structured and cohesive manner. The phases are clearly defined in the Makefile, enabling efficient configuration, execution and reproducibility of tasks. Figure 1 visually depicts the workflow, illustrating the logical progression of data and processes from preparation to final analysis.

**Figure 1.**
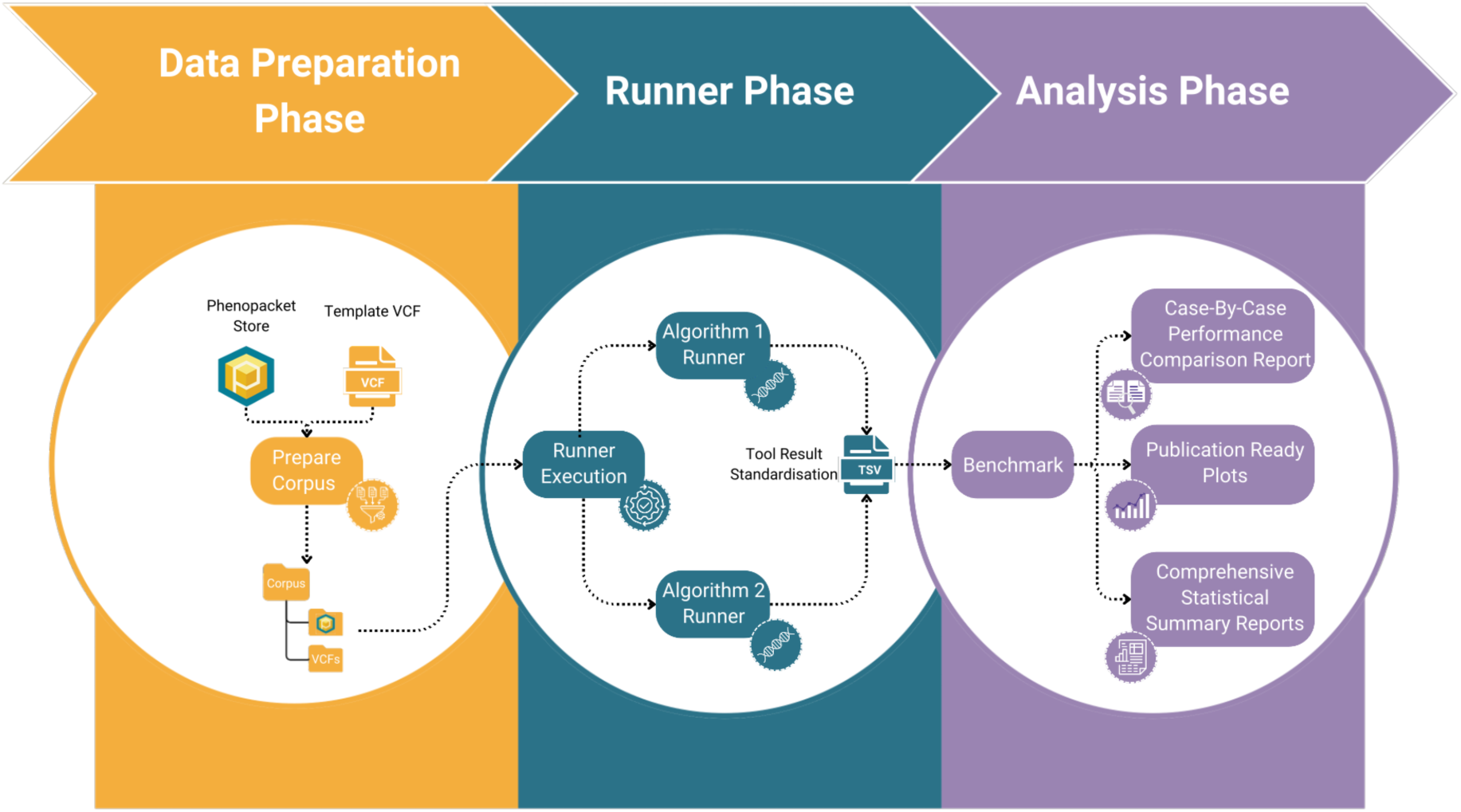
Pipeline workflow illustrative description from an experimental setting where two algorithms are compared. The workflow starts with the preparation of corpora which are then consumed by the concrete runners. Lastly, a consolidated report evaluating the results of the experiments is generated.

### Test corpora in PhEval

#### 4K corpus

Our main test corpus is the “phenopacket-store” [22] by Danis et al. The corpus at the time of this writing comprised 4916 GA4GH phenopackets (0.1.12) representing 277 diseases and 2872 unique pathogenic alleles, curated from 605 publications. Each phenopacket includes: a set of HPO terms describing the phenotypic profiles and a diagnosis, encompassing comprehensive information about the individuals previously documented in published case reports. This collection is the first large-scale, standardised set of case-level phenotypic information derived from detailed clinical data in literature case reports.

#### LIRICAL corpus

A small comparison corpus created for benchmarking the LIRICAL system [10] which contains 385 case reports. Variant information was generated by spiking causal genetic variants into a whole exome hg19 VCF file sourced from a healthy patient from the Genome in a Bottle dataset [23].

#### Synthetic corpus based on HPOA

We also provide a corpus of 8,245 synthetically generated patients produced with *phenotype2phenopacket*. The synthetic corpus is created from the HPO annotations provided by the Monarch Initiative, specifically using the 2024-04-26 release. The primary objective of this corpus is to simulate patient profiles specifically tailored to a specific disease. The corpus construction involves two steps, designed to represent the phenotypic characteristics associated with the disease while including noise.

HPO provides information on all possible phenotypes associated with that disease and the frequency at which they occur; we refer to this as the HPO term’s frequency value. In the first step of the corpus generation, a subset of HPO phenotypic terms is randomly selected for the disease, with the selection size varying between 20% and 75% of the total available terms. Each term undergoes scrutiny based on a randomly generated frequency value. If this value falls below the annotated frequency found in the HPO database, the term is deemed suitable for inclusion in the patient profile, to ensure diversity, terms lacking an annotated frequency are assigned a random frequency ranging from 0.25 to 0.75. This step also considers age onsets, where the patient’s age is generated within the range specified by the onset criteria from the HPO age of onset annotations. To introduce further variability, up to 33% of the total selected terms are supplemented with entirely random terms that are added independently of the frequency evaluation process.

Following the initial term selection, the profile is then further refined. In the next step, a subset of the selected terms is subjected to adjustments aimed at increasing or decreasing specificity. Each selected term undergoes a random number of adjustments within the ontology tree, involving movements up or down the tree by a specified number of nodes, ranging from 1-5. These adjustment steps are constrained to prevent terms from ascending beyond the top-level term “Abnormality of the X” in the HPO hierarchy. This adjustment process enriches the patient profiles with variability and specificity, aligning them with the complexity of real-world clinical scenarios.

To validate that the synthetic corpus maintains the phenotypic characteristics associated with the disease while introducing variability, we compared three independently generated sets of patient profiles and the 4K corpus to the disease models (Figure 2). This comparison evaluates how well the synthetic profiles reflect the phenotypical characteristics of the disease models and demonstrates that the synthetic profiles exhibit a similar distribution of similarity scores to the 4K corpus, providing evidence that the key phenotypic features of the disease are preserved.

**Figure 2.**
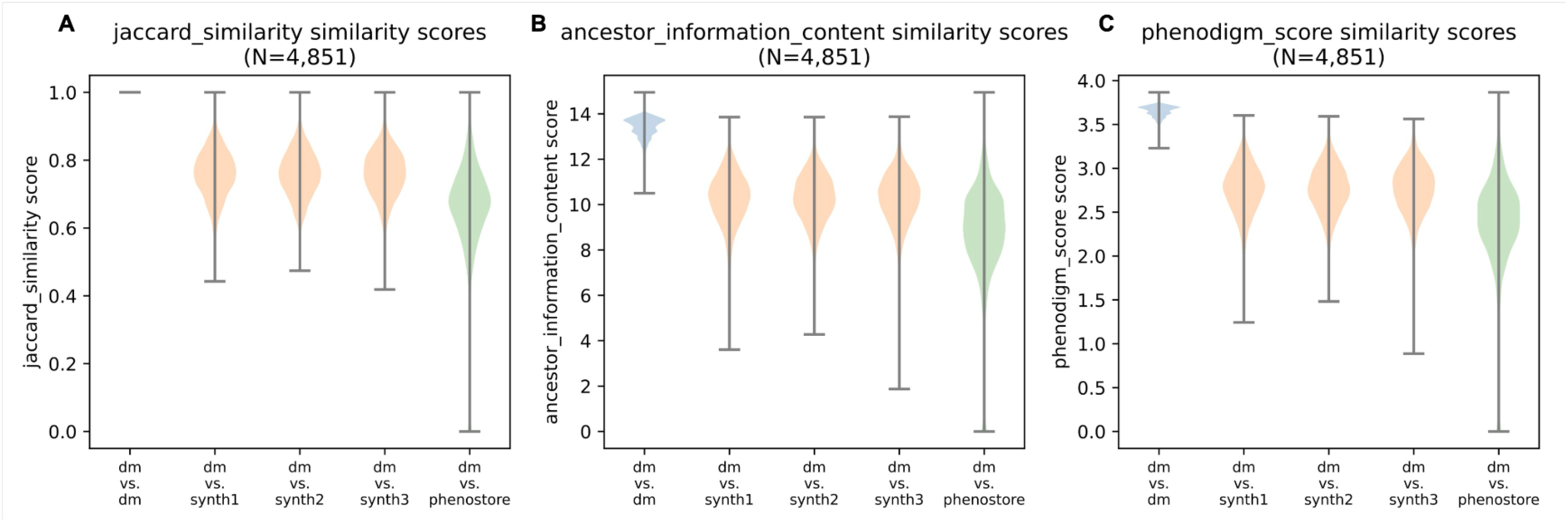
Violin plots showing similarity scores across three metrics: Jaccard similarity (A), ancestor information content (B), and Phenodigm scores (C). The plots compare disease models (dm) with three independently generated sets of synthetic patient profiles (synth1, synth2, synth3) and the 4K corpus.

Across all three metrics - Jaccard similarity, which measures term overlap; ancestor information content, which assesses phenotypic specificity within the ontology; and Phenodigm scores, which evaluates overall phenotypic alignment - the synthetic profiles show distributions that align closely with the 4K corpus. The violin plots depict that the synthetic profiles consistently achieve high similarity scores relative to the disease models, comparable to the spread observed for the 4K corpus. This indicates that the synthetic corpus retains disease-relevant phenotypic characteristics while introducing a degree of variability that mirrors the diversity inherent in real-world clinical data. Additionally, the spread in similarity scores highlights the ability of the synthetic corpus to capture phenotypic nuances across different diseases.

#### Structural variants corpus

We provide GA4GH phenopackets which are used to represent 188 structural variants known to be associated with specific diseases, extracted from 182 case reports published in 146 scientific articles [24]. Structural variant information is generated by spiking causal genetic structural variants into a whole exome hg38 structural variant VCF from a healthy individual from the Genome in a Bottle dataset.

#### Phen2Gene corpus

We provide 281 curated phenopackets using the primary data in the Phen2Gene benchmarking study [11]. This corpus represents individuals who were diagnosed with single-gene diseases and contain detailed phenotypic profiles as well as the known disease-causing gene.

#### PhEval Corpus Scramble Utility

PhEval provides a “scramble” utility designed to randomise the individual phenotypic profile within an existing phenopacket. The term “scrambling” refers to the introduction of noise. This functionality is particularly valuable for assessing algorithm performance for prioritising variants and genes in the presence of noise. By applying different scramble factors on a corpus, the changes in performance can be observed. This approach stands in contrast to the synthetic corpus based on HPOA which generates a new phenopacket based on a disease and statistical information about phenotypic distribution of that disease. The scramble factor, specified in the range of 0-1, determines the extent of scrambling, with 0 indicating no scrambling and 1 indicating a complete randomisation of the phenotypic profile. The scrambling process uses a proportional approach for the generation of test data, leveraging existing patient data from a pre-existing corpus. It works as follows. Let **s** be a scrambling factor between 0 and 1 and let a given patient’s phenopacket have **n** phenotypic terms. The scrambling is performed by randomly selecting **s * n** of the phenotypic terms from the profile and modifying them, while leaving the unselected terms unmodified. The modification is as follows: **s*n/2** or half of the selected terms are replaced by their parent term in the ontology, while the other half of the selected terms are replaced by another term in the ontology (with no concern for the original term). During the process of scrambling, some terms are retained, others are converted to parent terms to decrease specificity, and additional random terms are introduced. Specifically, if a scramble factor of 0.5 was employed, half of the total number original phenotypic terms would be retained, regardless of the depth of the original term being replaced; a quarter of the total number would be converted to parent terms and the remaining quarter would be random terms added to the profile. This proportion based scrambling strategy accommodates the diverse lengths of phenotypic profiles, allowing researchers to systematically explore algorithm robustness under varied conditions.

## Results

In the following, we describe the outcomes of a benchmarking process conducted using the latest available versions of each VGPA at the time of evaluation, with the default recommended parameter settings applied to each tool. The evaluation was performed on the 4K corpus, with pre-processing steps applied, using the PhEval *prepare-corpus* command that included removing phenopackets with missing gene and variant fields, converting gene identifiers to the ENSEMBL namespace and spiking the relevant variant into a template exome VCF. The benchmarking comparisons include: (1) Exomiser (run in phenotype only mode), Phen2Gene, PhenoGenius [25], and GADO [26], (2) Exomiser (run with VCF files), LIRICAL, and AI-MARRVEL [27], and (3) Exomiser and SvAnna [24] for structural variant analysis. We have selected this set of configurations to effectively illustrate the capabilities of PhEval, not to perform a comprehensive empirical study which would comprise dozens configurations. Therefore, for brevity, we omit details about the implemented tools and their configurations here but these are available from the GitHub repository as described below.

The benchmarking analyses revealed key distinctions in the performance of the evaluated phenotype-driven VGPAs across the different comparisons tested (Figure 3). Exomiser consistently demonstrated stronger diagnostic performance across all evaluations. In analyses restricted to phenotype-based inputs alone (Figure 3A), it achieved the highest proportion of causative genes ranked within the top candidates, with a notably higher MRR compared to GADO, Phen2Gene, and PhenoGenius. ROC (Figure 3B) and precision-recall (Figure 3C) curves demonstrated comparable sensitivity and specificity across all tools, with AUC values for ROC consistently high, while precision-recall AUC were lower across the board.

**Figure 3:**
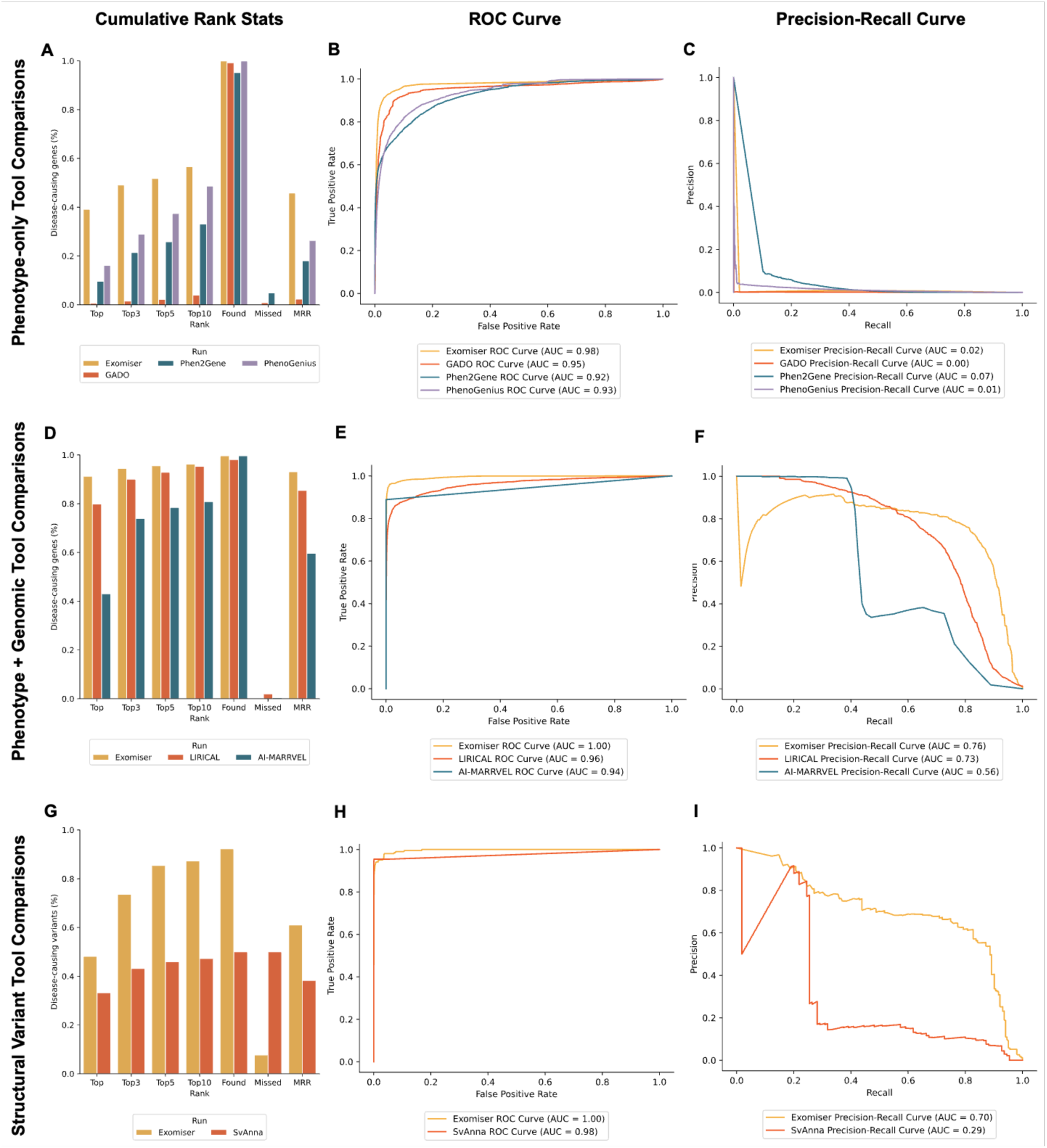
Performance comparison of phenotype-driven prioritisation tools across three analysis types: phenotype-only and phenotype-plus-genomic analyses, both conducted on the 4K corpus, and structural variant analyses conducted on the structural variant corpus. Rank-based metrics (A, D, G) show the percentage of known entities ranked within the top 1, 3, 5 and 10 ranked candidates along with the percentage of found and missed entities. MRR scores, ranging from 0 to 1, are also presented, with higher values indicating better ranking performance. ROC curves (B, E, H) and Precision Recall curves (C, F, I) depict statistical methods based on the confusion matrix of the causative entities.

When integrating phenotypic and genotypic data, Exomiser continued to exhibit stronger performance in comparison to the other tools in the evaluation, capturing more causative genes in higher ranks (Figure 3D). Compared to using phenotype-based inputs alone, Exomiser identified a greater number of highly ranked candidates. This improvement in ranking is reflected in the precision-recall analyses, where the integration of genomic data increased the AUC from 0.02 to 0.76 (Figure 3F). LIRICAL also performed robustly, maintaining a strong balance between highly-ranked candidates and recall, achieving performance comparable to Exomiser for top 10 candidates. By contrast, AI-MARRVEL, while capable of identifying causative genes, captured fewer in the top ranked positions compared to Exomiser and LIRICAL. Its precision also declined more rapidly as recall increased, reflecting limitations in maintaining precision across broader ranking thresholds.

For structural variant analysis, Exomiser outperformed SvAnna by identifying a higher proportion of causative variants and ranking them at higher positions (Figure 3G). While SvAnna demonstrated reasonable effectiveness in broader rankings, its precision and recall decreased at higher thresholds, highlighting the complexities of accurately prioritising structural variants (Figure 3I). This benchmarking process highlights the value of PhEval in facilitating standardised, reproducible evaluations of VGPAs, providing critical insights into tool-specific trade-offs and diagnostic capabilities.

A complete breakdown of the generated metrics can be found in Supplementary Table 1 for phenotype-only comparisons, Supplementary Table 2 for phenotype and genomic comparisons, and Supplementary Table 3 for structural variant comparisons.

## Discussion

Existing benchmarks predominantly report recall-based metrics, often measuring the algorithm’s capability to capture all relevant candidates in a reasonable set of ranked candidates. For example, the Phen2Gene benchmark assessed the performance of Phen2Gene against three other gene prioritisation tools. In this benchmark, researchers compared the number of prioritised genes that were found in the top 10, 50, 100 and 250 ranks for solved cases [11]. The concentration on recall metrics only provides a partial view of the algorithm’s performance and may inadvertently encourage algorithms to generate longer lists of equally ranked candidates, increasing the chances of identifying crucial genes or variants in the top X hits. However, a recall-centric approach can also lead to a higher rate of false positives (low precision), which may not always align with the demands of diverse research contexts, e.g., under-resourced diagnostic laboratories that can only properly interpret a handful of variants per case.

This perspective highlights the need for a balanced evaluation that takes both precision and recall into account when evaluating VGPAs for diagnostic use; as well as other metrics which may aid in the fine-tuning of algorithm processes. As reported in our results, we have provided a comprehensive evaluation using both ranking and binary statistics to demonstrate the performance of different configurations.There are existing benchmarks that go beyond the confines of recall metrics and provide a better picture into algorithm performance. For example, other efforts have explored insights into the sensitivity and specificity of algorithms by using area under the curve (AUC) from Receiver Operating Characteristic (ROC) in addition to an algorithm’s precision [12, 28, 29]. However, it can be difficult to draw direct comparisons between benchmarks that report different metrics due to the variations in the level of detail provided by these insights highlighting the need for the standardisation of an evaluation process and furnishing a consistent set of metrics and evaluation protocols.

PhEval was designed to provide a transparent, easy to use experimental framework that addresses issues such as the one illustrated by the anecdote above. A fully transparent and executable pipeline in a known data orchestration format (GNU-make) and standardised runner configurations, and versioned and standardised test data are key features that increase the transparency and reproducibility of VGPA benchmarks.

Standardising the benchmarking process of phenotype-driven VGPAs is complicated further by the increasing need of leveraging phenotype data alongside more classic gene-focused methods [30]. Modern VGPAs increasingly rely on phenotype data to improve diagnostic yield. An example of a VGPA tool that leverages cross-species phenotype data is Exomiser. Exomiser has also played a pivotal role in numerous projects and pipelines dedicated to novel gene discovery. Leveraging organism phenotype data proves to be an important step in the context of functional validation, as demonstrated by Pippucci et al., who used Exomiser to enhance the prioritisation of candidate genes in a case of epileptic encephalopathy, ultimately identifying a previously undiscovered mutation in CACNA2D2 to be causative of the disease [31]. PhenomeNET Variant Predictor (PVP), an alternative variant prioritisation algorithm, has also shown that the incorporation of mouse and fish phenotype data is especially useful in instances where human phenotypic information for a specific gene is lacking. Notably, PVP found substantial improvements in variant ranking when incorporating organism phenotype data in comparison to human alone. In a specific example genomic information relating to Hypotrichosis 8 was screened by PVP, initially variant rs766783183 present in the gene KRT25 was ranked at 172 (without the inclusion of model organism data); when this data was included this variant’s prioritisation ranking significantly improved to rank 8, this variant was ultimately the confirmed molecular diagnosis for Hypotrichosis 8 [32]. These cases demonstrate the importance of assessing VGPA performance based on their ability to integrate phenotypic data from model organisms, underscoring the need for a standardised evaluation framework.

Systematic benchmarking of VGPAs is important to monitor diagnostic yield in an environment that involves complex interactions between phenotype data and algorithms, but no standardised frameworks exist that support the entire benchmarking lifecycle. The closest one is VPMBench [33] which automates the benchmarking of variant prioritisation algorithms, but not the benchmarking of VGPAs that specifically leverage phenotype data. As phenotype-driven VGPAs integrate diverse phenotypic data sources, they require a distinct evaluation framework that addresses the added complexity in both algorithm design and evaluation. Leveraging phenotype data not only increases the complexity of the algorithms and therefore the importance of systematic benchmarking to monitor performance over time. It also increases the complexity of the evaluation itself, because of additional requirements on test data, test data pre-processing and analysis. PhEval has been designed to standardise the evaluation process for VGPAs with a specific focus on algorithms that leverage phenotype data. As we can see in Figure 3 the standardisation of analytical results supported by rigorous statistical methods, enables straightforward comparisons among various VGPAs. Individual phenopackets correspond to case descriptions that contain critical information that can inform the VGPA process. Every test in PhEval corresponds to a phenopacket, which not only ensures that every test case is appropriately standardised, but also that future test data that is already standardised as phenopackets can be seamlessly integrated into PhEval without complex transformation pipelines.

To facilitate wide uptake of the framework for experimental studies, an easy way to integrate existing VGPAs with often idiosyncratic distributions, configuration requirements and technology dependencies is needed. PhEval provides an easy-to-implement system to integrate any runner, which requires the implementation of a handful of methods; see Section on the Implementation of the PhEval CLI and VGPA runners. The instructions we provide for implementing runners such as these(https://monarch-initiative.github.io/pheval/developing_a_pheval_plugin/) also include a simple way to enable their publication on PyPi. This way, other experimenters can simply install runners that have already been implemented, like Exomiser, Phen2Gene and X, e.g. “pip install pheval.exomiser”.

### Limitations

#### Corpus bias

The most significant limitation of any specific framework for assessing the performance of diagnostic tools is the lack of publicly available real clinical data. This is no different in the case of PhEval. In practice, we execute PhEval on a number of private corpora such as the rare disease component of the 100,000 Genomes Project in the Genomics England (GEL) research environment, diagnosed cases from the Deciphering Developmental Disorders (DDD) project [34], and a retinal cohort [35].

The lack of a gold standard complicates comparative analyses among different algorithms. Variant prioritisation algorithms typically depend on curated databases of known disease-associated variants including specific subsets categorised by their established clinical significance, such as pathogenic or benign variants. These databases of curated disease-gene or disease-variant relationships, for example HPO and ClinVar, are often used for benchmarking [6, 36]. This dependence introduces a potential source of data circularity, as many cases in the 4K corpus were extracted from scientific literature, which may overlap with ClinVar pathogenic variants used in VGPA training. Even though these overlaps may occur, these are limited to a small subset of our corpus and ClinVar. A perfect evaluation corpus requires disease sequencing programs to hold back novel clinical diagnoses from ClinVar as in the recent Critical Assessment of Genomic Interpretation (CAGI) rare genomes project challenge [37]. However, resources and patient consent complications limit the size of such datasets (the number of solved cases in the CAGI challenge was only 35) and they cannot be shared openly which reduces their utility for systematic benchmarking.While these datasets serve as valuable resources for benchmarking, they are limited in scope and may not fully represent the genetic variability and diseases encountered in real-world scenarios. Specifically, the phenotype data typically presents as a merge of all possible phenotypes for a disease (HPOA) or a disease label (ClinVar) resulting in a loss of specific individual phenotype information that could otherwise be linked to genetic variants. In light of these challenges, it becomes imperative for the research community to work collectively toward establishing standardised benchmarking methodologies and datasets to advance the ability to openly and rigorously evaluate and compare VGPAs.

While the development of a proper representative gold standard corpus of real clinical samples is still largely out of sight, we are constantly working on increasing our public test corpora. We are currently building a phenopackets store (https://github.com/monarch-initiative/phenopacket-store), which contains an extensive collection of Phenopackets that represent real individuals with Mendelian disease reported in case reports in the literature.

#### Tool integration challenges

While the study highlights PhEval’s compatibility with a range of VGPAs, the inclusion of additional tools was limited by both maintenance and technical constraints. Several tools could not be added as plugins and evaluated due to issues with installation often stemming due to outdated documentation, lack of programmatic access, or infrequent updates, with some tools not being updated in the last 3 years. This lack of updates limits their clinical utility, as hundreds of new disease-gene associations are discovered annually, and diagnostics rely on tools that can query up-to-date knowledge. These factors constrained the scope of the benchmarking analysis. Future benchmarking efforts will depend on the continued development and maintenance of VGPAs to enable wider evaluations.

## Conclusions

Variant prioritisation is critical for diagnosis of rare and genetic conditions at scale. To effectively support diagnostics, VGPA methods increasingly need to leverage all the available data, in particular phenotype. As our ability to leverage phenotype data in more sophisticated ways increases, for example by including gene-to-phenotype associations from across different species, the complexity of the system increases as well. To ensure accurate evaluations of VGPA tools and monitoring their performance across versions, robust methods must be developed to evaluate the complex interplay between algorithms and data. PhEval is the first framework that takes phenotype data directly into account during the VGPA benchmarking process by standardising input data as GA4GH phenopackets.

**Table 1:** VGPA tools evaluated as part of the experiment.

|  | Corpus | VGPA Tool | Version |
| --- | --- | --- | --- |
| Phenotype-only Analysis | 4K corpus | Exomiser | 14.0.2 & 2406 db release |
|  |  | GADO | 1.0.4 |
|  |  | Phen2Gene | 1.2.3 |
|  |  | PhenoGenius | 1.1.1 |
| Phenotype + Genomic Analysis (phenotypes + VCF) | 4K corpus | Exomiser | 14.0.2 & 2406 db release |
|  |  | LIRICAL | 2.0.2 |
|  |  | AI-MARRVEL | 0.1.0 |
| Phenotype + Structural Variant Analysis | Structural variant corpus | Exomiser | 14.0.2 & 2406 db release |
|  |  | SvAnna | 1.0.4 |

The entire experimental setup, with the exception of the private corpora, is available at

## Availability and requirements

Project name: PhEval

Project home page: https://github.com/monarch-initiative/pheval

Operating system(s): UNIX

Programming language: Python

Other requirements: Docker, Python libraries

Licence: Apache License 2.0

Any restrictions to use by non-academics: none

## List of abbreviations

NGS: Next-generation sequencing
WES: Whole-exome sequencing
WGS: Whole-genome sequencing
VGPA: Variant and gene prioritisation algorithm
HPO: Human phenotype ontology
HGMD: Human gene mutation database
PyPi: Python package index
CLI: Command line interface
MRR: Mean Reciprocal Rank
NDCG: Normalised discounted cumulative gain
ROC: Receiver operator characteristic
AUC: Area under the curve
PVP: PhenomeNET variant predictor
LLM: Large language model
GEL: Genomics england
DDD: Deciphering developmental disorders

## Declarations

### Ethics approval and consent to participate

Not applicable

### Consent for publication

Not applicable

### Availability of data and materials

The package source code repository can be accessed via GitHub at https://github.com/monarch-initiative/pheval. The documentation can be found at https://monarch-initiative.github.io/pheval/. The data used to produce the results in the study can be found at https://zenodo.org/records/14679713 and the code used for reproducing the benchmarking from this paper is available at https://github.com/monarch-initiative/pheval-paper.

The PhEval corpora are under constant development and can be found at https://github.com/monarch-initiative/pheval/tree/main/corpora.

### Authors’ contributions

YB developed the PhEval framework and runners, and authored the manuscript. VdS developed the experimental pipelines and authored the manuscript; KGC, AO tested the pipeline and authored the manuscript; MH, NLH, DRK, NMM, AO edited the manuscript, NM, JAM, CJM, DS and JJ coordinated the project and provided input on the design; DOS and PNR provided expert feedback on the design and corpus processing. All authors have read and approved the manuscript.

## Acknowledgements

Not applicable.

## References

1. Groft SC, Posada M, Taruscio D: Progress, challenges and global approaches to rare diseases. Acta Paediatr. 2021, 110:2711–2716.

2. Jia J, Shi T: Towards efficiency in rare disease research: what is distinctive and important? Sci. China Life Sci. 2017, 60:686–691.

3. Kruse J, Mueller R, Aghdassi AA, Lerch MM, Salloch S: Genetic Testing for Rare Diseases: A Systematic Review of Ethical Aspects. Front. Genet. 2021, 12:701988.

4. The 1000 Genomes Project Consortium: A global reference for human genetic variation. Nature 2015, 526:68.

5. Single Nucleotide Polymorphism [10.1016/B978-0-12-812537-3.00012-3].

6. Gargano MA, Matentzoglu N, Coleman B, et al.: The Human Phenotype Ontology in 2024: phenotypes around the world. Nucleic Acids Res. 2024, 52:D1333–D1346.

7. Foreman J, Brent S, Perrett D, Bevan AP, Hunt SE, Cunningham F, Hurles ME, Firth HV: DECIPHER: Supporting the interpretation and sharing of rare disease phenotype-linked variant data to advance diagnosis and research. Hum. Mutat. 2022, 43:682–697.

8. Jacobsen JOB, Kelly C, Cipriani V, Genomics England Research Consortium, Mungall CJ, Reese J, Danis D, Robinson PN, Smedley D: Phenotype-driven approaches to enhance variant prioritization and diagnosis of rare disease. Hum. Mutat. 2022, 43:1071.

9. Smedley D, Jacobsen JOB, Jäger M, Köhler S, Holtgrewe M, Schubach M, Siragusa E, Zemojtel T, Buske OJ, Washington NL, Bone WP, Haendel MA, Robinson PN: Next-generation diagnostics and disease-gene discovery with the Exomiser. Nat. Protoc. 2015, 10:2004–2015.

10. Robinson PN, Ravanmehr V, Jacobsen JOB, Danis D, Zhang XA, Carmody LC, Gargano MA, Thaxton CL, UNC Biocuration Core, Karlebach G, Reese J, Holtgrewe M, Köhler S, McMurry JA, Haendel MA, Smedley D: Interpretable Clinical Genomics with a Likelihood Ratio Paradigm. Am. J. Hum. Genet. 2020, 107:403–417.

11. Zhao M, Havrilla JM, Fang L, Chen Y, Peng J, Liu C, Wu C, Sarmady M, Botas P, Isla J, Lyon GJ, Weng C, Wang K: Phen2Gene: rapid phenotype-driven gene prioritization for rare diseases. NAR Genom Bioinform 2020, 2:lqaa032.

12. Robinson PN, Köhler S, Oellrich A, Sanger Mouse Genetics Project, Wang K, Mungall CJ, Lewis SE, Washington N, Bauer S, Seelow D, Krawitz P, Gilissen C, Haendel M, Smedley D: Improved exome prioritization of disease genes through cross-species phenotype comparison. Genome Res. 2014, 24:340–348.

13. Thompson R, Papakonstantinou Ntalis A, Beltran S, Töpf A, de Paula Estephan E, Polavarapu K, ’t Hoen PAC, Missier P, Lochmüller H: Increasing phenotypic annotation improves the diagnostic rate of exome sequencing in a rare neuromuscular disorder. Hum. Mutat. 2019, 40:1797–1812.

14. Putman TE, Schaper K, Matentzoglu N, Rubinetti VP, Alquaddoomi FS, Cox C, Caufield JH, Elsarboukh G, Gehrke S, Hegde H, Reese JT, Braun I, Bruskiewich RM, Cappelletti L, Carbon S, Caron AR, Chan LE, Chute CG, Cortes KG, De Souza V, Fontana T, Harris NL, Hartley EL, Hurwitz E, Jacobsen JOB, Krishnamurthy M, Laraway BJ, McLaughlin JA, McMurry JA, Moxon SAT, Mullen KR, O’Neil ST, Shefchek KA, Stefancsik R, Toro S, Vasilevsky NA, Walls RL, Whetzel PL, Osumi-Sutherland D, Smedley D, Robinson PN, Mungall CJ, Haendel MA, Munoz-Torres MC: The Monarch Initiative in 2024: an analytic platform integrating phenotypes, genes and diseases across species. Nucleic Acids Res. 2024, 52.

15. Bone WP, Washington NL, Buske OJ, Adams DR, Davis J, Draper D, Flynn ED, Girdea M, Godfrey R, Golas G, Groden C, Jacobsen J, Köhler S, Lee EMJ, Links AE, Markello TC, Mungall CJ, Nehrebecky M, Robinson PN, Sincan M, Soldatos AG, Tifft CJ, Toro C, Trang H, Valkanas E, Vasilevsky N, Wahl C, Wolfe LA, Boerkoel CF, Brudno M, Haendel MA, Gahl WA, Smedley D: Computational evaluation of exome sequence data using human and model organism phenotypes improves diagnostic efficiency. Genet. Med. 2015, 18:608–617.

16. Peng C, Dieck S, Schmid A, Ahmad A, Knaus A, Wenzel M, Mehnert L, Zirn B, Haack T, Ossowski S, Wagner M, Brunet T, Ehmke N, Danyel M, Rosnev S, Kamphans T, Nadav G, Fleischer N, Fröhlich H, Krawitz P: CADA: phenotype-driven gene prioritization based on a case-enriched knowledge graph. NAR Genom Bioinform 2021, 3:lqab078.

17. Pengelly RJ, Alom T, Zhang Z, Hunt D, Ennis S, Collins A: Evaluating phenotype-driven approaches for genetic diagnoses from exomes in a clinical setting. Sci. Rep. 2017, 7:1–7.

18. Ebiki M, Okazaki T, Kai M, Adachi K, Nanba E: Comparison of Causative Variant Prioritization Tools Using Next-generation Sequencing Data in Japanese Patients with Mendelian Disorders. Yonago Acta Med. 2019, 62:244–252.

19. Yuan X, Wang J, Dai B, Sun Y, Zhang K, Chen F, Peng Q, Huang Y, Zhang X, Chen J, Xu X, Chuan J, Mu W, Li H, Fang P, Gong Q, Zhang P: Evaluation of phenotype-driven gene prioritization methods for Mendelian diseases. Brief. Bioinform. 2022, 23:bbac019.

20. Jacobsen JOB, Kelly C, Cipriani V, Robinson PN, Smedley D: Evaluation of phenotype-driven gene prioritization methods for Mendelian diseases. Brief. Bioinform. 2022, 23:bbac188.

21. Jacobsen JOB, Baudis M, Baynam GS, Beckmann JS, Beltran S, Buske OJ, Callahan TJ, Chute CG, Courtot M, Danis D, Elemento O, Essenwanger A, Freimuth RR, Gargano MA, Groza T, Hamosh A, Harris NL, Kaliyaperumal R, Lloyd KCK, Khalifa A, Krawitz PM, Köhler S, Laraway BJ, Lehväslaiho H, Matalonga L, McMurry JA, Metke-Jimenez A, Mungall CJ, Munoz-Torres MC, Ogishima S, Papakonstantinou A, Piscia D, Pontikos N, Queralt-Rosinach N, Roos M, Sass J, Schofield PN, Seelow D, Siapos A, Smedley D, Smith LD, Steinhaus R, Sundaramurthi JC, Swietlik EM, Thun S, Vasilevsky NA, Wagner AH, Warner JL, Weiland C, GAGH Phenopacket Modeling Consortium, Haendel MA, Robinson PN: The GA4GH Phenopacket schema defines a computable representation of clinical data. Nat. Biotechnol. 2022, 40:817–820.

22. Danis D, Bamshad MJ, Bridges Y, Cacheiro P, Carmody LC, Chong JX, Coleman B, Dalgleish R, Freeman PJ, Graefe ASL, Groza T, Jacobsen JOB, Klocperk A, Kusters M, Ladewig MS, Marcello AJ, Mattina T, Mungall CJ, Munoz-Torres MC, Reese JT, Rehburg F, Reis BCS, Schuetz C, Smedley D, Strauss T, Sundaramurthi JC, Thun S, Wissink K, Wagstaff JF, Zocche D, Haendel MA, Robinson PN: A corpus of GA4GH Phenopackets: case-level phenotyping for genomic diagnostics and discovery. bioRxiv 2024.

23. Zook JM, Catoe D, McDaniel J, Vang L, Spies N, Sidow A, Weng Z, Liu Y, Mason CE, Alexander N, Henaff E, McIntyre ABR, Chandramohan D, Chen F, Jaeger E, Moshrefi A, Pham K, Stedman W, Liang T, Saghbini M, Dzakula Z, Hastie A, Cao H, Deikus G, Schadt E, Sebra R, Bashir A, Truty RM, Chang CC, Gulbahce N, Zhao K, Ghosh S, Hyland F, Fu Y, Chaisson M, Xiao C, Trow J, Sherry ST, Zaranek AW, Ball M, Bobe J, Estep P, Church GM, Marks P, Kyriazopoulou-Panagiotopoulou S, Zheng GXY, Schnall-Levin M, Ordonez HS, Mudivarti PA, Giorda K, Sheng Y, Rypdal KB, Salit M: Extensive sequencing of seven human genomes to characterize benchmark reference materials. Sci Data 2016, 3:160025.

24. Danis D, Jacobsen JOB, Balachandran P, Zhu Q, Yilmaz F, Reese J, Haimel M, Lyon GJ, Helbig I, Mungall CJ, Beck CR, Lee C, Smedley D, Robinson PN: SvAnna: efficient and accurate pathogenicity prediction of coding and regulatory structural variants in long-read genome sequencing. Genome Med. 2022, 14:44.

25. Yauy K, Duforet-Frebourg N, Testard Q, Beaumeunier S, Audoux J, Simard B, Larue D, Blum MGB, Bernard V, Genevieve D, Bertrand D, PhenoGenius consortium, Philippe N, Thevenon J: Learning phenotypic patterns in genetic diseases by symptom interaction modeling. medRxiv 2022:2022.07.29.22278181.

26. Deelen P, van Dam S, Herkert JC, Karjalainen JM, Brugge H, Abbott KM, van Diemen CC, van der Zwaag PA, Gerkes EH, Zonneveld-Huijssoon E, Boer-Bergsma JJ, Folkertsma P, Gillett T, van der Velde KJ, Kanninga R, van den Akker PC, Jan SZ, Hoorntje ET, te Rijdt WP, Vos YJ, Jongbloed JDH, van Ravenswaaij-Arts CMA, Sinke R, Sikkema-Raddatz B, Kerstjens-Frederikse WS, Swertz MA, Franke L: Improving the diagnostic yield of exome-sequencing by predicting gene–phenotype associations using large-scale gene expression analysis. Nature Communications 2019, 10:1–13.

27. Mao D, Liu C, Wang L, Ai-Ouran R, Deisseroth C, Pasupuleti S, Kim SY, Li L, Rosenfeld JA, Meng L, Burrage LC, Wangler MF, Yamamoto S, Undiagnosed Diseases Network, Santana M, Perez V, Shukla P, Eng CM, Lee B, Yuan B, Xia F, Bellen HJ, Liu P, Liu Z: AI-MARRVEL - A Knowledge-Driven AI System for Diagnosing Mendelian Disorders. NEJM AI 2024, 1.

28. Danzi MC, Dohrn MF, Fazal S, Beijer D, Rebelo AP, Cintra V, Züchner S: Deep structured learning for variant prioritization in Mendelian diseases. Nat. Commun. 2023, 14:4167.

29. Requena T, Gallego-Martinez A, Lopez-Escamez JA: A pipeline combining multiple strategies for prioritizing heterozygous variants for the identification of candidate genes in exome datasets. Hum. Genomics 2017, 11:11.

30. Seaby EG, Rehm HL, O’Donnell-Luria A: Strategies to Uplift Novel Mendelian Gene Discovery for Improved Clinical Outcomes. Front. Genet. 2021, 12.

31. Pippucci T, Parmeggiani A, Palombo F, Maresca A, Angius A, Crisponi L, Cucca F, Liguori R, Valentino ML, Seri M, Carelli V: A Novel Null Homozygous Mutation Confirms CACNA2D2 as a Gene Mutated in Epileptic Encephalopathy. PLoS One 2013, 8.

32. Boudellioua I, Mahamad Razali RB, Kulmanov M, Hashish Y, Bajic VB, Goncalves-Serra E, Schoenmakers N, Gkoutos GV, Schofield PN, Hoehndorf R: Semantic prioritization of novel causative genomic variants. PLoS Comput. Biol. 2017, 13:e1005500.

33. Ruscheinski A, Reimler AL, Ewald R, Uhrmacher AM: VPMBench: a test bench for variant prioritization methods. BMC Bioinformatics 2021, 22:1–15.

34. Wright CF, Fitzgerald TW, Jones WD, Clayton S, McRae JF, van Kogelenberg M, King DA, Ambridge K, Barrett DM, Bayzetinova T, Bevan AP, Bragin E, Chatzimichali EA, Gribble S, Jones P, Krishnappa N, Mason LE, Miller R, Morley KI, Parthiban V, Prigmore E, Rajan D, Sifrim A, Swaminathan GJ, Tivey AR, Middleton A, Parker M, Carter NP, Barrett JC, Hurles ME, FitzPatrick DR, Firth HV, DDD study: Genetic diagnosis of developmental disorders in the DDD study: a scalable analysis of genome-wide research data. Lancet 2015, 385:1305–1314.

35. Kelly C, Szabo A, Pontikos N, Arno G, Robinson PN, Jacobsen JOB, Smedley D, Cipriani V: Phenotype-aware prioritisation of rare Mendelian disease variants. Trends Genet. 2022.

36. Landrum MJ, Lee JM, Benson M, Brown GR, Chao C, Chitipiralla S, Gu B, Hart J, Hoffman D, Jang W, Karapetyan K, Katz K, Liu C, Maddipatla Z, Malheiro A, McDaniel K, Ovetsky M, Riley G, Zhou G, Holmes JB, Kattman BL, Maglott DR: ClinVar: improving access to variant interpretations and supporting evidence. Nucleic Acids Res. 2018, 46:D1062–D1067.

37. Stenton SL, O’Leary MC, Lemire G, VanNoy GE, DiTroia S, Ganesh VS, Groopman E, O’Heir E, Mangilog B, Osei-Owusu I, Pais LS, Serrano J, Singer-Berk M, Weisburd B, Wilson MW, Austin-Tse C, Abdelhakim M, Althagafi A, Babbi G, Bellazzi R, Bovo S, Carta MG, Casadio R, Coenen P-J, De Paoli F, Floris M, Gajapathy M, Hoehndorf R, Jacobsen JOB, Joseph T, Kamandula A, Katsonis P, Kint C, Lichtarge O, Limongelli I, Lu Y, Magni P, Mamidi TKK, Martelli PL, Mulargia M, Nicora G, Nykamp K, Pejaver V, Peng Y, Pham THC, Podda MS, Rao A, Rizzo E, Saipradeep VG, Savojardo C, Schols P, Shen Y, Sivadasan N, Smedley D, Soru D, Srinivasan R, Sun Y, Sunderam U, Tan W, Tiwari N, Wang X, Wang Y, Williams A, Worthey EA, Yin R, You Y, Zeiberg D, Zucca S, Bakolitsa C, Brenner SE, Fullerton SM, Radivojac P, Rehm HL, O’Donnell-Luria A: Critical assessment of variant prioritization methods for rare disease diagnosis within the rare genomes project. Hum Genomics 2024, 18:44.

