## Supplementary Table 1 for "Towards a standard benchmark for phenotype-driven variant and gene prioritisation algorithms: PhEval - Phenotypic inference Evaluation framework"

Supplementary Table 1. Ranking and binary classification statistics reported for the phenotype-only analysis tested on the 4K corpus.

| Statistic | Exomiser | GADO | Phen2Gene | PhenoGenius |
| --- | --- | --- | --- | --- |
| Top | 1828 | 29 | 449 | 756 |
| Top 3 | 2295 | 70 | 1002 | 1355 |
| Top 5 | 2422 | 102 | 1208 | 1751 |
| Top 10 | 2646 | 184 | 1548 | 2275 |
| Found | 4674 | 4641 | 4452 | 4674 |
| Total | 4677 | 4677 | 4677 | 4677 |
| Mean Reciprocal Rank | 0.458 | 0.023 | 0.180 | 0.263 |
| Precision@1 | 0.391 | 0.006 | 0.096 | 0.162 |
| Precision@3 | 0.164 | 0.005 | 0.071 | 0.097 |
| Precision@5 | 0.104 | 0.004 | 0.052 | 0.075 |
| Precision@10 | 0.057 | 0.004 | 0.033 | 0.049 |
| MAP@1 | 0.391 | 0.006 | 0.096 | 0.162 |
| MAP@3 | 0.436 | 0.010 | 0.150 | 0.217 |
| MAP@5 | 0.442 | 0.012 | 0.160 | 0.237 |
| MAP@10 | 0.448 | 0.014 | 0.169 | 0.252 |
| F-beta score@1 | 0.391 | 0.006 | 0.096 | 0.162 |
| F-beta score@3 | 0.231 | 0.006 | 0.082 | 0.121 |
| F-beta score@5 | 0.164 | 0.005 | 0.067 | 0.102 |
| F-beta score@10 | 0.099 | 0.005 | 0.049 | 0.075 |
| NDCG@3 | 0.260 | 0.015 | 0.151 | 0.136 |
| NDCG@5 | 0.249 | 0.016 | 0.163 | 0.165 |
| NDCG@10 | 0.221 | 0.022 | 0.172 | 0.184 |
| Sensitivity | 0.391 | 0.006 | 0.096 | 0.162 |
| Specificity | 1.000 | 1.000 | 1.000 | 1.000 |
| Precision | 0.455 | 0.006 | 0.098 | 0.163 |
| Negative predictive value | 1.000 | 1.000 | 1.000 | 1.000 |
| False positive rate | 0.000 | 0.000 | 0.000 | 0.000 |
| False discovery rate | 0.545 | 0.994 | 0.902 | 0.837 |
| False negative rate | 0.609 | 0.994 | 0.904 | 0.838 |
| Accuracy | 1.000 | 1.000 | 1.000 | 1.000 |
| F1 score | 0.421 | 0.006 | 0.097 | 0.162 |
| Matthew’s correlation coefficient | 0.422 | 0.006 | 0.097 | 0.162 |
