## Supplementary Table 2 for "Towards a standard benchmark for phenotype-driven variant and gene prioritisation algorithms: PhEval - Phenotypic inference Evaluation framework"

Supplementary Table 2. Ranking and binary classification statistics reported for the phenotype and genomic analysis tested on the 4K corpus.

| Statistic | Exomiser | LIRICAL | AI-MARRVEL |
| --- | --- | --- | --- |
| Top | 4097 | 3585 | 1931 |
| Top 3 | 4241 | 4042 | 3318 |
| Top 5 | 4290 | 4169 | 3522 |
| Top 10 | 4321 | 4282 | 3631 |
| Found | 4476 | 4404 | 4474 |
| Total | 4491 | 4491 | 4491 |
| Mean reciprocal rank | 0.931 | 0.854 | 0.596 |
| Precision@1 | 0.912 | 0.798 | 0.430 |
| Precision@3 | 0.315 | 0.300 | 0.246 |
| Precision@5 | 0.191 | 0.186 | 0.157 |
| Precision@10 | 0.096 | 0.095 | 0.081 |
| MAP@1 | 0.912 | 0.798 | 0.430 |
| MAP@3 | 0.926 | 0.843 | 0.580 |
| MAP@5 | 0.928 | 0.850 | 0.590 |
| MAP@10 | 0.929 | 0.853 | 0.594 |
| F beta score@1 | 0.912 | 0.798 | 0.430 |
| F beta score@3 | 0.468 | 0.436 | 0.313 |
| F beta score@5 | 0.316 | 0.301 | 0.230 |
| F beta score@10 | 0.174 | 0.170 | 0.136 |
| NDCG@3 | 0.529 | 0.508 | 0.406 |
| NDCG@5 | 0.467 | 0.466 | 0.375 |
| NDCG@10 | 0.381 | 0.394 | 0.322 |
| Sensitivity | 0.912 | 0.798 | 0.430 |
| Specificity | 1.000 | 0.998 | 1.000 |
| Precision | 0.932 | 0.800 | 0.439 |
| Negative predictive value | 1.000 | 0.998 | 1.000 |
| False positive rate | 0.000 | 0.002 | 0.000 |
| False discovery rate | 0.068 | 0.200 | 0.561 |
| False negative rate | 0.088 | 0.202 | 0.570 |
| Accuracy | 1.000 | 0.995 | 1.000 |
| F1 score | 0.922 | 0.799 | 0.434 |
| Matthew’s correlation coefficient | 0.922 | 0.797 | 0.434 |
