## Supplementary Table 3 for "Towards a standard benchmark for phenotype-driven variant and gene prioritisation algorithms: PhEval - Phenotypic inference Evaluation framework"

Supplementary Table 3. Ranking and binary classification statistics reported for the structural variant analysis tested on the structural variant corpus.

| Statistic | Exomiser | SvAnna |
| --- | --- | --- |
| Top | 106 | 73 |
| Top 3 | 162 | 95 |
| Top 5 | 188 | 101 |
| Top 10 | 192 | 104 |
| Found | 203 | 110 |
| Total | 220 | 220 |
| Mean reciprocal rank | 0.610 | 0.383 |
| Precision@1 | 0.482 | 0.332 |
| Precision@3 | 0.245 | 0.144 |
| Precision@5 | 0.171 | 0.092 |
| Precision@10 | 0.087 | 0.047 |
| MAP@1 | 0.486 | 0.336 |
| MAP@3 | 0.597 | 0.381 |
| MAP@5 | 0.633 | 0.388 |
| MAP@10 | 0.636 | 0.390 |
| F-beta score@1 | 0.482 | 0.332 |
| F-beta score@3 | 0.325 | 0.201 |
| F-beta score@5 | 0.252 | 0.144 |
| F-beta score@10 | 0.148 | 0.083 |
| NDCG@3 | 0.447 | 0.749 |
| NDCG@5 | 0.495 | 0.719 |
| NDCG@10 | 0.426 | 0.683 |
| Sensitivity | 0.482 | 0.333 |
| Specificity | 1.000 | 1.000 |
| Precision | 0.741 | 0.424 |
| Negative predictive value | 0.999 | 1.000 |
| False positive rate | 0.000 | 0.000 |
| False discovery rate | 0.259 | 0.576 |
| False negative rate | 0.518 | 0.667 |
| Accuracy | 0.998 | 1.000 |
| F1 score | 0.584 | 0.373 |
| Matthew’s correlation coefficient | 0.597 | 0.376 |
